# Characterization of Unique Pathological Features of COVID-Associated Coagulopathy: Studies with AC70 hACE2 Transgenic Mice Highly Permissive to SARS-CoV-2 Infection

**DOI:** 10.1101/2023.10.30.564680

**Authors:** Aleksandra K. Drelich, Kempaiah Rayavara, Jason Hsu, Panatda Saenkham-Huntsinger, Barbara M. Judy, Vivian Tat, Thomas G. Ksiazek, Bi-Hung Peng, Chien-Te K. Tseng

## Abstract

COVID-associated coagulopathy seemly plays a key role in post-acute sequelae of SARS-CoV-2 infection. However, the underlying pathophysiological mechanisms are poorly understood, largely due to the lack of suitable animal models that recapitulate key clinical and pathological symptoms. Here, we fully characterized AC70 line of human ACE2 transgenic (AC70 hACE2 Tg) mice for SARS-CoV-2 infection. We noted that this model is highly permissive to SARS-CoV-2 with values of 50% lethal dose and infectious dose as ∼ 3 and ∼ 0.5 TCID_50_ of SARS-CoV-2, respectively. Mice infected with 10^5^ TCID_50_ of SARS-CoV-2 rapidly succumbed to infection with 100% mortality within 5 days. Lung and brain were the prime tissues harboring high viral titers, accompanied by histopathology. However, viral RNA and many inflammatory mediators could be readily detectable in other organs, suggesting the nature of a systemic infection. Lethal challenge of AC70 hACE2 Tg mice caused acute onset of leukopenia, lymphopenia, along with an increased neutrophil-to-lymphocyte ratio. Importantly, infected animals recapitulated key features of COVID-19-associated coagulopathy, including significantly elevated levels of D-dimer, t-PA, PAI-1, and circulating NETs, along with activated platelet/endothelium marker. Immunohistochemical staining with anti-PF4 antibody revealed profound platelet aggregates especially within blocked veins of the lungs. ANXA2 is known to interact with S100A10 to form heterotetrametric complexes, serving as coreceptors for t-PA to regulate membrane fibrinolysis. Thus, our results revealing elevated IgG type anti-ANXA2 antibody production, downregulated *de novo* ANXA2/S100A10 synthesis, and reduced AnxA2/S100A10 association in infected mice support an important role of this protein in the pathogenesis of acute COVID-19. In summary, we showed that acute SARS-CoV-2 infection of AC70 hACE2 Tg mice triggered a hypercoagulable state coexisting with ill-regulated fibrinolysis, accompanied by dysregulation of ANXA2 system, which might serve as druggable targets for development of antithrombotic and/or anti-fibrinolytic agents to attenuate pathogenesis of COVID-19.

**Author Summary:** Accumulating evidence strongly suggests that COVID-associated coagulopathy characterized by dysregulation of the coagulation cascade, fibrinolysis system and pulmonary microvascular immune-thrombosis during different stages of SARS-CoV-2 infection may have a “yet-to-be fully defined” impact on the development of post-acute sequela of COVID-19. Herein we initially reported a comprehensively characterized AC70 hACE2 Tg mouse model for SARS-CoV-2 infection and disease. We next demonstrated the subsequent onset of imbalanced coagulation and fibrinolysis pathways in infected Tg mice, focusing on dysregulated formation of ANXA2/S100A10 complexes, key coreceptors for t-PA that regulates membrane fibrinolysis, in which elevated production of autoantibodies against ANXA2 induced by SARS-CoV-2 might play an intriguing role. Taken together, we demonstrated that AC70 hACE2 Tg mice lethally challenged with SARS-CoV-2 recapitulated several features of COVID-associated coagulopathy observed in patients and highlighted the potential role of ANXA2 in this phenomenon. Thus, ANXA2 might serve as a potentially novel druggable target to attenuate COVID-19-associated thrombotic events.

## Introduction

Over 770 million COVID-19 cases have been reported worldwide, with more than 100 million cases in the US alone, resulting in over 6.9 and 1.1 million deaths, respectively [World Health Organization, WHO COVID-19 disease Dashboard, 2020]. While the speedy interventions and widespread usage of effective vaccines and drugs, including neutralizing monoclonal antibodies (mAbs), immunomodulators, and antivirals (e.g., Paxlovid or Lagevrio), have greatly mitigated the pandemic threat worldwide, newly emerged SARS-CoV-2 variants-of-concern (VOCs) capable of escaping neutralizing mAbs, such as Omicron BA.2, XBB.1.5, and other sub-lineages, make it a great challenge to fully contain COVID-19 worldwide.

Many of the early studies on COVID-19 were largely focused on severe or even fatal cases upon acute SARS-CoV-2 infection without paying much attention to those that experienced lasting various symptoms of COVID-19. New reports show that ∼30% of patients treated for COVID-19 developed post-acute sequela of COVID-19 (PASC), also known as Long-COVID, including fatigue, shortness of breath, migraines, loss of smell or taste, anxiety, or depression, even after clinical recovery [1].

Although the exact underlying pathophysiological mechanism(s) of PASC remains elusive, it seems to be very complex in nature which might be initiated upon ill-regulated innate immune responses that failed to control the primary viral infection, resulting in the onset of exacerbated inflammatory responses, platelet aggregation, neutrophil extracellular trap (NET) formation, endothelium activation, and even promptly provoking autoantibody production [2, 3]. The subsequent progression of these processes could eventually lead to structural and functional impairment of multiple organs, particularly the lung, brain, heart, and kidney. Current evidence suggests that an ongoing process of thrombi formation and/or persistent micro-immuno-thrombosis may be a key player in the development of the main physical post-acute sequelae of COVID-19 [4].

Under normal physiological conditions, the steady state of hemostasis is sustained by the dynamic balance between blood clot formation, via activating a cascade of clotting factors, and the canonical fibrinolytic pathway initiated by tissue plasminogen activator (t-PA) released predominantly from activated endothelial cells (ECs) that activates plasminogen to its active form, plasmin, enabling the degradation of fibrin mesh. Thus, acute SARS-CoV-2 infection in some COVID-19 patients might lead to impairment of this balance by inducing alteration in both processes. Indeed, markedly elevated levels of D-dimers, the final products of fibrin degradation, were frequently associated with patients suffering from SARS-CoV-2 infection and was correlated with disease severity [5], suggesting hyper-coagulation and hyper-fibrinolytic state during the disease process. Conversely, the elevated expression of plasminogen activator inhibitor 1 (PAI-1) in severe COVID patients might indicate a paradoxical coexistence of hypo-fibrinolysis as well [6]. Post-mortem samples collected from patients that succumbed to SARS-CoV-2 infection often showed deposition of fibrin at sites of small vessels and capillaries, and extracellular spaces, further indicating aberrant activation of the fibrinolysis cascade [7]. Importantly, many reports have indicated that thrombosis frequently occurred in severe COVID-19 patients, despite the prophylactical and/or therapeutic treatments with anticoagulants [8, 9], thereby suggesting that multiple pathways might be involved to synergistically create a prothrombotic milieu at various stages of active SARS-CoV-2 infection; however, the exact mechanisms of action are not clear thus far.

Although the relevance of the coagulopathy has not been fully clarified, the existence of readily detectable levels of antiphospholipid antibody (aPL) is commonly observed in COVID-19 patients. For instances, up to 50% of COVID-19 patients have been shown to possess elevated levels of lupus anticoagulants, a class of immunoglobulins specifically targeting phospholipid binding proteins [10]. A recent study demonstrated that one of such autoantibodies targeted Annexin A2 (ANXA2) was readily elevated among fatal cases, compared to those being hospitalized but not severe COVID-19 patients [11]. ANXA2 is a calcium-dependent phospholipid-binding protein that exists as a monomer within the cytoplasm or forms a heterotertrameric complexes (AIIt) localized at the extracellular membranes with two S100A10 (p11) small molecules acting as a co-receptor for t-PA, tightly involved in cell surface fibrinolysis [12]. Additionally, ANXA2 has been implicated in the pathogenesis of many pathogens, including SARS-CoV [13]. Thus, an altered ANXA2-mediated fibrinolysis, in part due to enhanced induction of autoantibodies against ANXA2 triggered upon SARS-CoV-2 infection, might contribute to the complex pathophysiological mechanisms of COVID-associated coagulopathy.

COVID-associated coagulopathy represents a major area of COVID-19 research to not only delineate its pathophysiological mechanisms but also develop effective countermeasures to protect against long-term consequences of coagulation disorders. Therefore, identifying an animal species permissive to SARS-CoV-2 infection and diseases that could faithfully recapitulate key features of coagulopathy is desirable to facilitate basic research on this phenomenon and test the preclinical efficacy of either repurposing or new drug interventions. Like SARS-CoV, SARS-CoV-2 uses human angiotensin-converting enzyme 2 (hACE2) as a functional receptor to enter permissive hosts for establishing its infection [14]. As we have generated and fully characterized several lineages of transgenic mice constitutively and systemically expressing hACE2 gene at varying intensities for SARS-CoV infection [15], in this study we initially fully characterized the response of AC70 hACE2 Tg mice to respiratory SARS-CoV-2 (US_WA1/2020) infection, with regard to kinetics and tissue distribution of viral replication, inflammatory responses, tissue histopathology, and the values of 50% lethal dose (LD_50_) and infectious dose (ID_50_), followed by assessing key features of coagulopathy associated with COVID frequently observed in severe human cases. We also demonstrated a dysregulated ANXA2 mechanism triggered by acute SARS-CoV-2 infection, that might serve as a potential druggable target for attenuating thrombotic events in PASC.

## Results

### Comprehensive characterization of AC70 hACE2 Tg mice against SARS-CoV-2 infection

Among several lineages of transgenic mice globally expressing hACE2 that we have established during the SARS-CoV era, the AC70 lineage of hACE2 Tg mice was best characterized as a lethal model of SARS-CoV infection and disease [15, 16]. As SARS-CoV-2 uses the same hACE2 as the entry receptor to the permissive hosts, we initially challenged AC70 hACE2 Tg mice through intranasal (i.n.) route with a high-dose (∼10^5^ TCID_50_ in 60 μl) of SARS-CoV-2 (US-WA1/2020) for assessing their permissiveness to infection. As shown in **Figs 1A and B**, we found that challenged mice suffered from a rapid onset of weight loss, other signs of illness (i.e., ruffled fur, lethargy, hunched posture, orbital tightening, reluctance to move when stimulated), and ultimately succumbed to infection within five days post infection (d.p.i.) with 100% mortality. As expected, their transgene-negative (Tg^-^) littermates were not permissive to SARS-CoV-2, as evidenced by the absence of morbidity (i.e., weight loss) and mortality.

**Fig 1.**
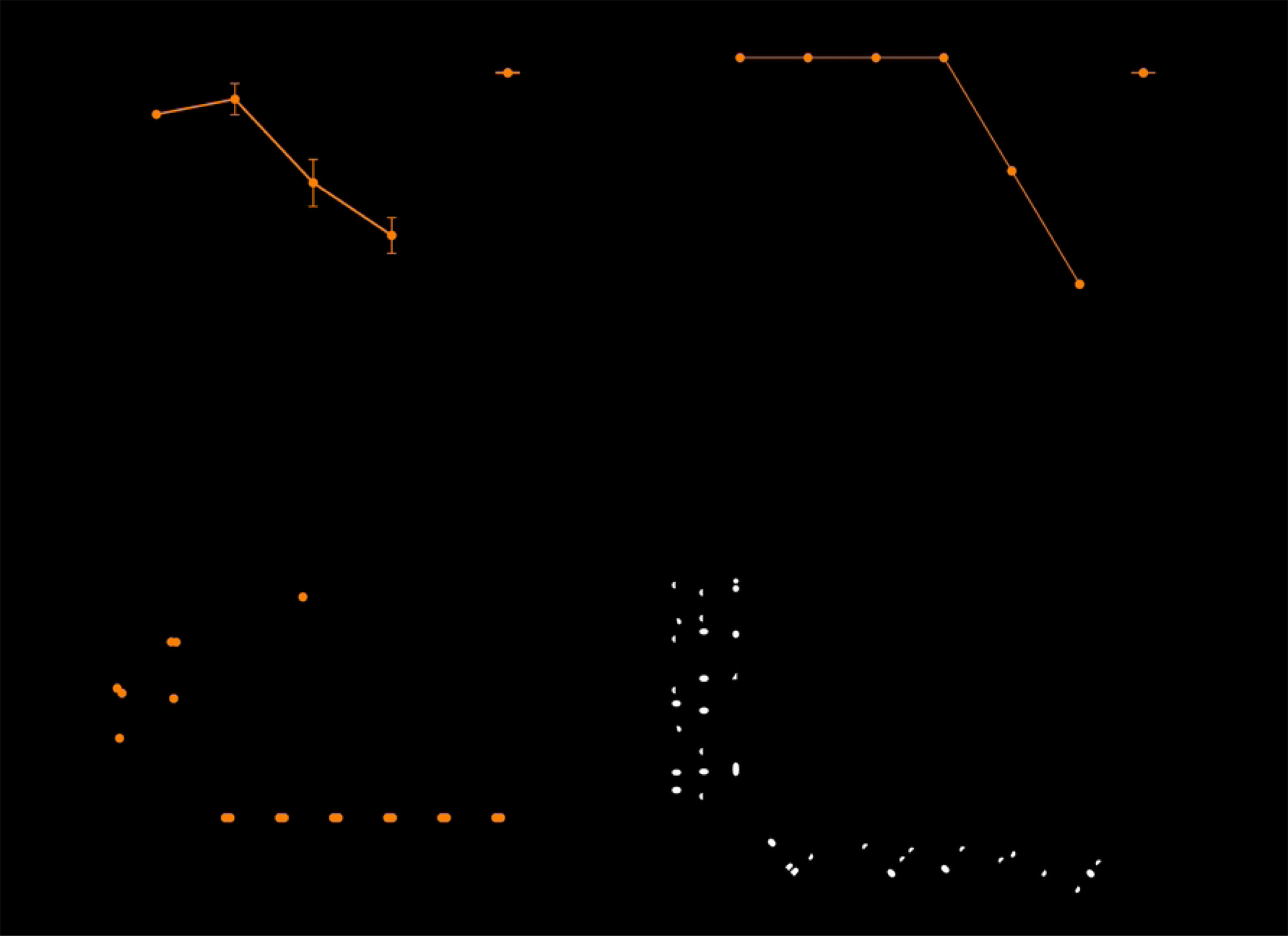
AC70 hACE2 transgene-positive (Tg^+^) mice challenged with a high dose of SARS-CoV-2, but not their transgene-negative (Tg^-^) littermates, resulted in an acute onset of morbidity and mortality. AC70 hACE2 (Tg⁺) and their Tg^-^ littermates (age-/sex-matched) were intranasally (i.n.) challenged with 10^5^ TCID_50_ of SARS-CoV-2. Challenged mice were monitored daily for morbidity (weight loss and other clinical signs) and mortality. (A) Average percentage (%) of daily body weight changes after viral infection, expressed as Mean ± SEM (standard error mean) were plotted. (B) Accumulative mortality rates of infected ACE2 Tg^+^ and Tg^-^ mice. Tissues/organs were harvested from infected Tg^+^ or Tg-mice at 4 d.p.i. (C) Yields of infectious progeny virus in various tissues harvested at 4 d.p.i. from AC70 Tg^+^ and Tg^-^ mice, respectively, were determined by using the standard Vero E6 cell-based infectivity assay and expressed as log_10_ TCID_50_ virus per gram of tissue. Limit of detection was ∼1.69 log_10_ TCID_50_/g. (D) Quantitative RT-PCR assays targeting SARS-CoV-2 E gene using total RNAs extracted from tissues collected were used to semi-quantify viral RNA. After normalizing to 18S RNA, relative fold changes in the transcripts of SARS-CoV-2 E gene expression in AC70 hACE2 Tg^+^ versus Tg^-^ mice were plotted.

We also explored viral yields and tissue distribution of SARS-CoV-2 in AC70 hACE2 Tg versus Tg^-^ mice at 4 d.p.i by using the standard Vero E6-based infectivity assay and qPCR targeting E gene of the virus. Among eight different tissues/organs assessed, i.e., brain, lung, liver, spleen, kidneys, heart, and small and large gastrointestinal (GI) tracts; lungs and brains were the only tissues/organs where infectious progeny virus at titers of 5.42 (lung) and 7.53 (brain) log_10_ TCID_50_/per gram of tissues were readily recovered from infected AC70 hACE2 Tg mice (**Fig 1C**). Consistent with the absence of readily noticeable onset of morbidity and mortality, we were unable to recover any infectious virus in all tissues/organs obtained from challenged AC70 Tg^-^ littermates. Whereas live viruses could only be recovered from the lung and brain, data derived from qPCR-based assays unambiguously indicated that respiratory viral challenge of AC70 hACE2 Tg mice resulted in a disseminated infection into other tissues/organs tested at various extents, with the highest levels of viral RNAs detected within lungs and brains (**Fig 1D**).

For assessing the host responses to productive SARS-CoV-2 infection, we investigated histopathology and inflammatory responses in AC70 hACE2 Tg mice at 4 d.p.i. We noted the existence of few foci of perivascular cuffing of infiltrating mononuclear cells within the lungs and areas of congestion/thrombosis (**Fig 2A**). Despite intense viral replication in the brain, small aggregates of mononuclear cells, along with microglial activation, could only be occasionally detected around arterioles within the striatum and brainstem (**Fig 2B**), compared to the same area of brain derived from AC70 Tg^-^ mice (**Fig 2C**). Additionally, we found few small foci of inflammatory infiltrates in the left ventricular wall of the heart with eosinophilic myocytes (**Fig 2D**). While cellular infiltrates could not be found in the spleen, liver, kidney, and GI tracts, we noticed tissue necrosis in the liver and small GI tracts, as shown in **Fig 2E and 2F**, respectively.

**Fig 2.**
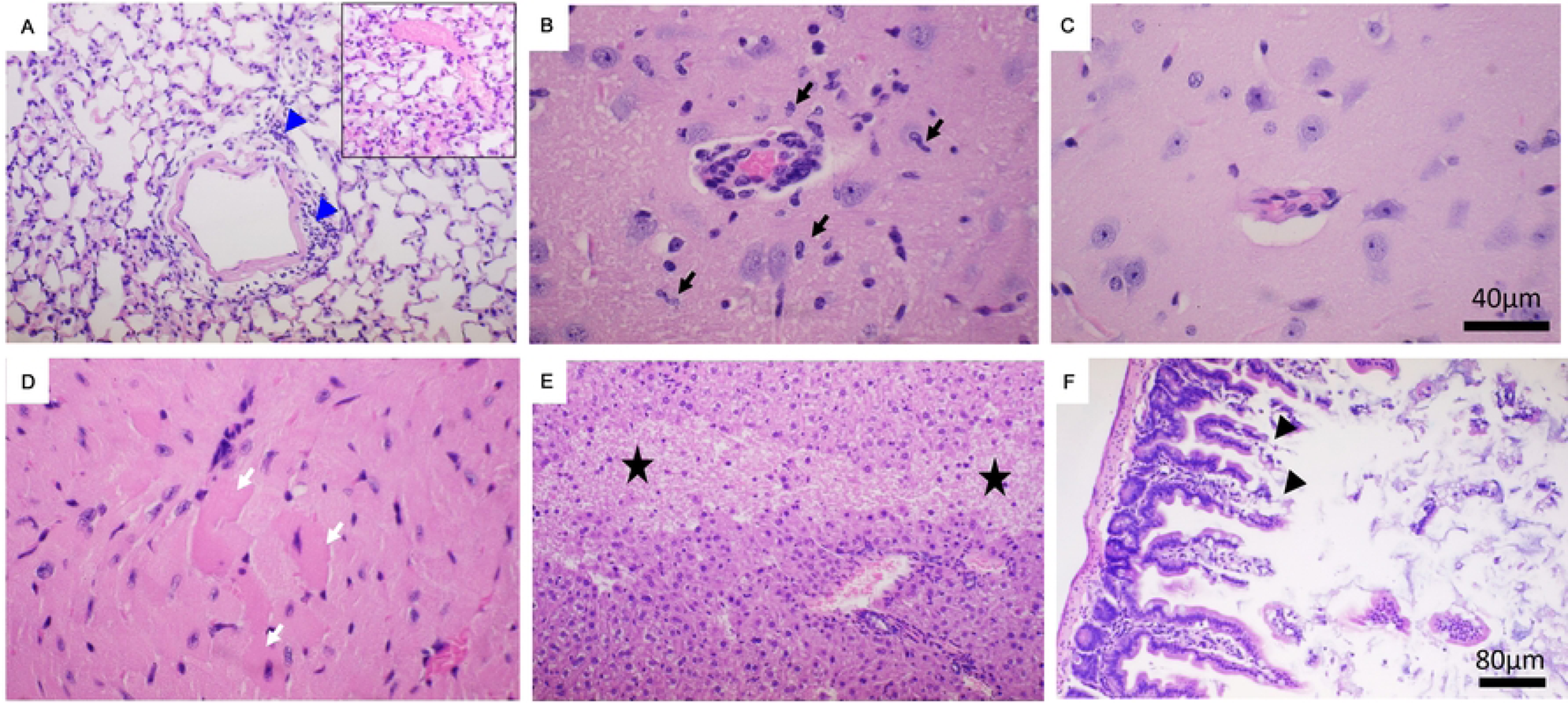
Histopathological examination of SARS-CoV-2-infected AC70 hACE2 Tg^+^ mice showed inflammatory infiltrates and tissue damage to varying extents. Various tissue specimens harvested at 4 d.p.i. from AC70 hACE2 Tg^+^ mice challenged with 10^5^ TCID_50_ of SARS-CoV-2 were subjected to the histopathology study. Briefly, paraffin-embedded, and formalin-fixed tissue sections were stained with hematoxylin and eosin (H&E), as described in Materials and Methods. Mild perivascular infiltrations were seen in the lung (**A, blue arrowheads**) with areas of congestion/thrombosis (**inset**). In the brain (**B**), peri-vascular infiltration and microglial activation (**B, black arrows**) was seen in infected AC70 hACE2 Tg^+^ mice compared to a similar area of their Tg^-^ littermates (**C**). Small foci of infiltrates in the heart with eosinophilic myocytes (**D, white arrows**). Tissue necrosis was seen in the liver (**E, black stars**) and the small intestine (**F, black arrowheads**). Note: Bar= 80µm for A, E and F; Bar= 40µm for B, C and D.

The revelation of tissue damage and inflammatory infiltrates in productively infected AC70 hACE2 Tg mice prompted us to quantify the cytokine and chemokine responses among tissues by using qPCR-based analysis. As shown in **Fig S1**, induction of type I IFNs was readily revealed within various tissues. Specifically, transcriptional levels of IFN-α were significantly increased in most organs tested, except livers. Transcription of IFN-β was upregulated in brains, spleens, and kidneys, whereas an enhanced transcription of IFN-γ transcription was detected in spleens and livers. Interestingly, transcription of IFN-ls was profoundly induced especially within the lungs and brains. Transcripts of other inflammatory mediators were induced predominantly within the lungs (i.e., CXCL10, IL-8, RANTES, MCP1, MIP1β and MX1), brains (i.e., CXCL-1, CXCL-10, IL-6, IL-8, TNF-α, RANTES and MX1), kidneys (i.e., CXCL-10 and RANTES) and spleens (i.e., IL-1β, IL-6, IL-10, and TNF-α).

Finally, we determined the values of 50% lethal dose (LD_50_) and infectious dose (ID_50_) of SARS-CoV-2 in our AC70 hACE2 Tg mice to better characterize it as a consistent preclinical animal model for studies of viral pathogenesis and development of effective medical countermeasures (MCMs) by using the same strategy we previously described [17]. As shown in **Fig S2** and **Table S1**, the values of LD_50_ and ID_50_ were ∼3 and ∼0.5 TCID50 of SARS-CoV-2, respectively.

Having well-characterized this AC70 hACE2 Tg mouse lineage with regards to permissiveness to SARS-CoV-2 infection and disease, we subsequently investigated whether some key pathophysiological features observed in patients with confirmed COVID-associated coagulopathy could be revealed in AC70 hACE2 Tg mice challenged with 10^5^ TCID_50_ of SARS-CoV-2. We explored whether hematological and coagulation/fibrinolysis disorders, few key indicators of a still growing list of COVID symptoms, could be observed in these lethally challenged Tg mice.

### Lethally challenged AC70 hACE2 Tg mice exhibited acute onsets of hematological alterations

Both lymphopenia and increased neutrophil-to-lymphocyte ratios (NLR) have been implicated as hallmarks to differentiate patients suffering from either mild or severe COVID-19 disease [18, 19]. To investigate if such hematological alterations could occur in AC70 hACE2 Tg mice, a group of 10 mice challenged (i.n.) with 10^5^ TCID_50_ of SARS-CoV-2 were analyzed. Briefly, peripheral blood specimens of individual animals were collected from mice at 6 days prior and at 1, 2, or 3 days after lethal challenge with SARS-CoV-2 and subjected to hematological analyses using a VetScan analyzer (VSpro, ABAXIS). We noted that infected mice rapidly induced profound leukopenia, starting at 1 d.p.i. and persisting through 3 d.p.i., in which the total white blood cell (WBC) counts dropped from the mean value of 9.2 K/uL prior to infection to 6.2 (p=0.0179), 5.6 (p=0.0038) and 2.5 (p<0.0001) K/uL at 1, 2, and 3 d.p.i., respectively (**Fig 3A**). Among the subsets of WBC, the lymphocyte (LY) counts were the most affected that were drastically reduced from 7.1 K/uL before infection to 4.0 (p<0.0001), 1.6 (p<0.0001) and 0.8 (p<0.0001) K/uL at 1, 2, and 3 d.p.i., respectively (**Fig 3B**). We also noted that values of NLRs were slightly elevated from 0.2 prior to infection to 0.4 at 1 d.p.i. and significantly increased to 2.3 (p<0.0001) at both 2- and 3 d.p.i., respectively (**Fig 3D**). Neutrophil (NE) counts significantly increased by 1.84-fold (p=0.0157) at the midpoint and fell back to normal level at day 3 p.i. (**Fig 3C**). Total counts of other WBC subsets, including monocyte (MO), eosinophil (EO), and basophile (BA), were not significantly altered (**data not shown**). Erythrocyte profile revealed slight reduction in red blood cell (RBC) counts and hemoglobin (HGB) levels early in the infection and gradually increased later with mean counts of 8.6 M/uL (p=0,02) and 13.3 g/dL (p=0.005) at day 3 p.i., when compared to those measured before infection, i.e., 7.5 and 11.5 for RBC and HGB, respectively (**Fig 3E and F**). Additionally, we noted the levels of hematocrit (HCT) increased significantly over time after infection with a mean count of 31.6, 35.3 (p=0.015) and 37.7 (p=0.0009) % at 1, 2, and 3 d.p.i., respectively, when compared to 29.2% in uninfected mice (**Fig 3G**). Thrombocyte (PLT) counts showed a decreasing trend, albeit insignificant, at the early phase of infection, that gradually rebound to thrombocytosis at the peak of infection with a mean count of 952 K/uL (p<0.0001) at day 3 p.i., when compared to those measured before infection with a mean count of 484 K/uL (**Fig 3H**).

**Fig 3.**
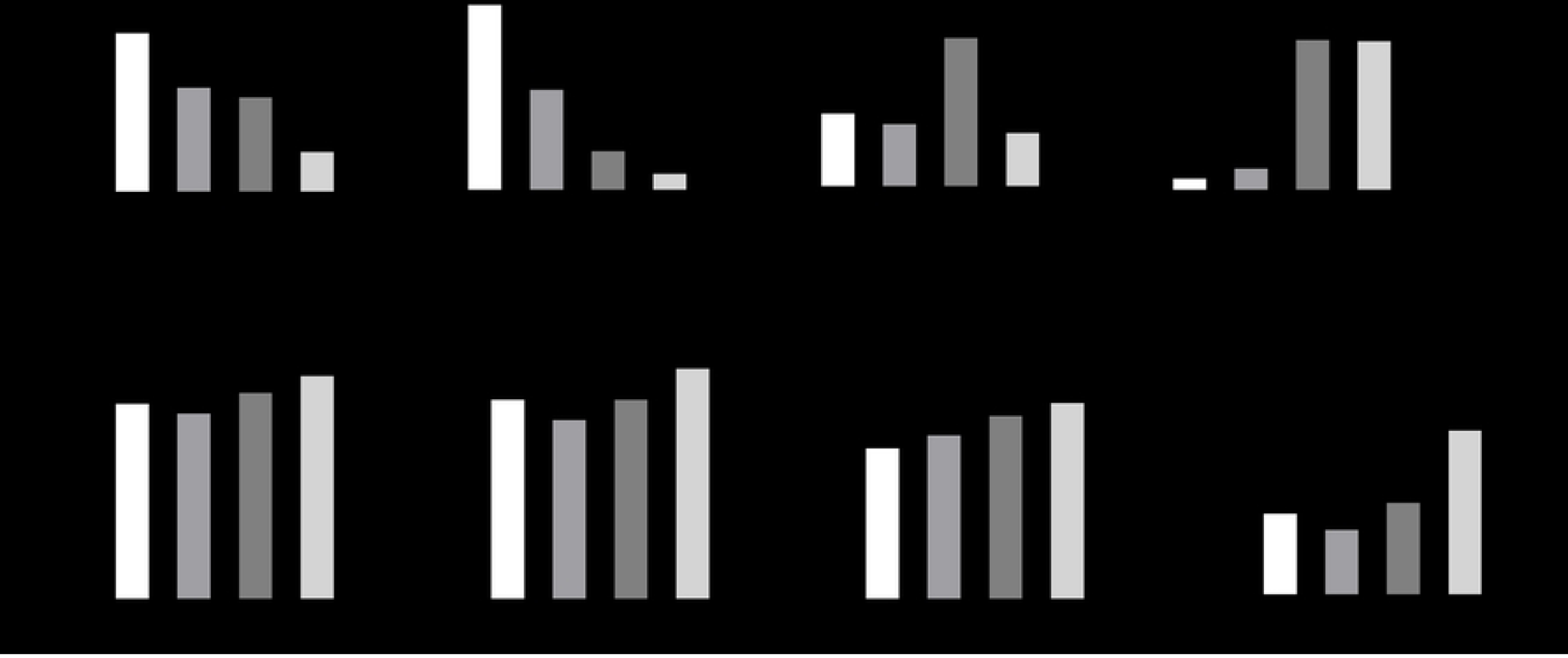
Lethally challenged AC70 hACE2 Tg mice exhibited profoundly altered hematology. White Blood Cell (WBC) counts (**A**), Lymphocyte (LY) counts (**B**), Neutrophil (NE) counts (**C**), The neutrophil-to-lymphocyte (NLR) ratio (**D**), Red Blood Cell (RBC) counts (**E**), Hemoglobin (HGB) levels (**F**), Hematocrit (HCT) levels (**G**), Platelet (PLT) counts (**H**). Data represent mean of 10 animals/time point. Statistical analyses between uninfected and infected mice at 1, 2, or 3 d.p.i. were performed using One-way ANOVA Multiple Comparison test. Results were considered significant with p < 0.05 (*p < 0.02; **p≤0.005, ***p=0.0009, ****p<0.0001).

### Lethal SARS-CoV-2 infection led to coagulation - fibrinolysis imbalance in AC70 hACE2 Tg mice

Since coagulation dysfunction plays a likely role in initiating the onset of multiple post-acute COVID sequelae, the assessment of expression of coagulation parameters has been commonly used as biomarkers for the rapid prediction of disease severity. Specifically, elevated plasma level of D-dimers, a fibrin degradation product (FDPs), has been frequently observed in COVID-19 patients with thrombotic complications and was associated with poor prognosis [20]. Therefore, we initially quantified circulating D-dimers in AC70 hACE2 Tg mice. Briefly, two groups of AC70 hACE2 Tg mice (n=5) were either challenged with 10^5^ TCID_50_ of SARS-CoV-2 or remained unchallenged, as controls. Both groups of mice were euthanized at 4 d.p.i., when infected mice manifested signs of severe disease. Plasma specimens collected prior to euthanasia were subjected to an ELISA-based analysis. As shown in **Fig 4A**, levels of D-dimers in the plasma were indeed significantly increased from the basal value of 2154 ng/mL prior to infection to 4057 ng/mL (p=0.0006), suggesting that dysregulated coagulation might have occurred in SARS-CoV-2-infected mice. As the fibrinolytic shutdown might coexist with hyper-coagulation in severe COVID-19 patients [21], we next quantified circulating plasminogen activator inhibitor 1 (PAI-1), a common marker of impaired endogenous fibrinolysis that is predominantly released from activated endothelial cells (ECs), in an attempt to explore if an impaired fibrinolytic pathway could be concomitantly induced upon SARS-CoV-2 infection. As shown in **Fig 4B**, the values of PAI-1 were markedly increased at 4 d.p.i. from the basal value of 204.4 (prior to infection) to 591.2 pg/mL at 4 dpi (p=0.0083). Despite the elevated PAI-1 expression, we also found that the levels of tissue-plasmin activator (t-PA) were increased from 36.3 (prior to infection) to 746.6 pg/mL at 4 dpi, an ∼18-folds increase (p < 0.0001), as shown in **Fig 4C**. Taken together, our data indicated that acute SARS-CoV-2 infection could concomitantly disturb the balance between coagulation and fibrinolytic pathways in AC70 hACE2 Tg mice.

**Fig 4.**
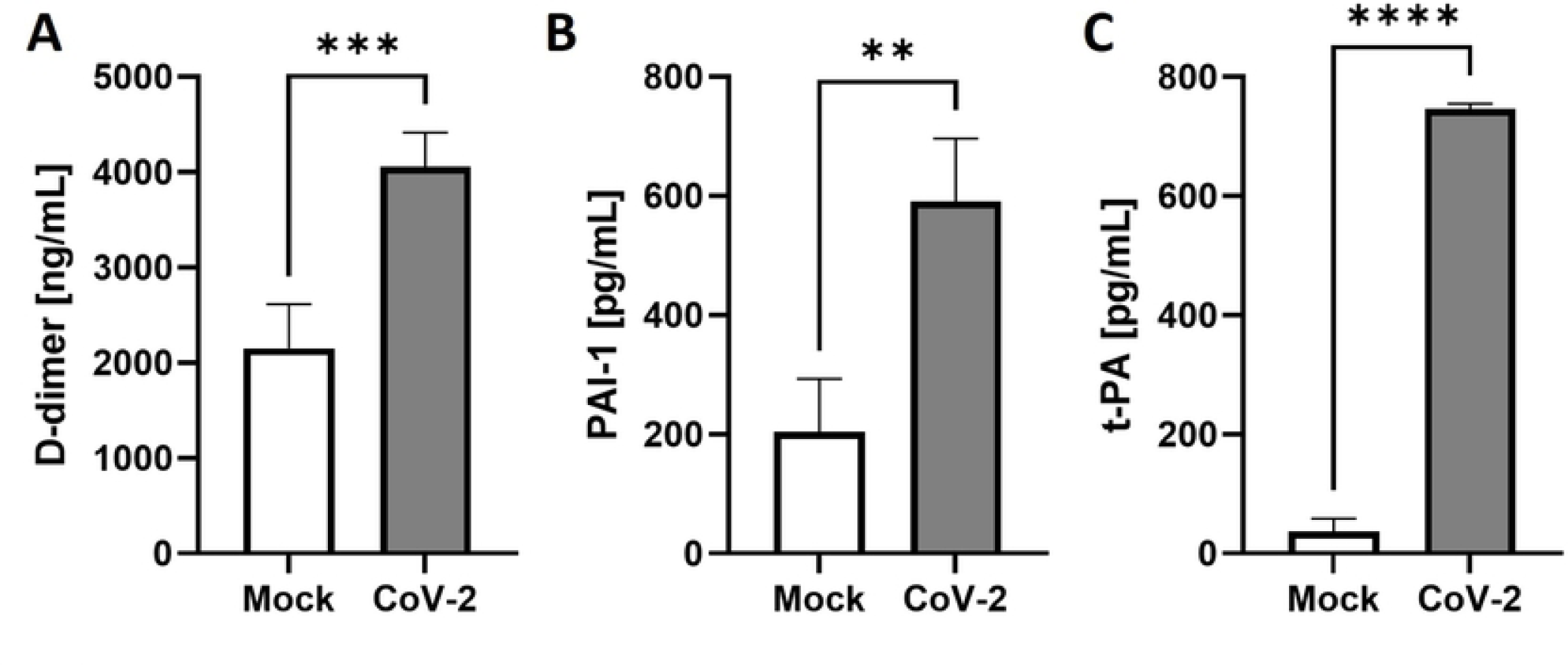
Lethal SARS-CoV-2 infection caused coagulation/ fibrinolysis abnormalities in AC70 hACE2 Tg mice. The concentration of D-dimers (**A**), Plasminogen Activator Inhibitor-1 (PAI-1) (**B**) and tissue Plasminogen Activator (t-PA) (**C**) in gamma irradiated EDTA-plasma of mock and SARS-CoV-2 infected mice (n=5 per group) were determined at 4 d.p.i. by ELISA. Results were considered significant with p < 0.05 (**p=0.0083, ***p=0.0006, ****p<0.0001).

### Lethal SARS-CoV-2 infection triggered both neutrophil and platelet activation in AC70 hACE2 Tg mice

Platelet hyperactivation and prothrombotic potential of Neutrophil Extracellular Traps (NETs) released from activated neutrophils are considered key aspects of the pathogenesis of COVID-associated coagulopathy [22–25]. Having revealed significantly altered coagulation system in infected mice, we next investigated whether signs of activated neutrophils/ NETs and platelets could occur in lethally infected AC70 hACE2 Tg mice. Plasma specimens collected at 4 d.p.i. were subjected to an ELISA-based analysis for quantifying the levels of myeloperoxidase (MPO) and soluble P-selectin, biomarkers of activated neutrophils/ NETs and platelets and/or endothelium, respectively. As shown in **Fig 5A**, a mean value of 12.4 ng/mL of circulating MPO was detected only in the specimens derived from infected mice, compared to the undetected levels (<0.61) in uninfected animals (p=0.037). We also noted the levels of P-selectin in the plasma were significantly increased to the level of 6332 pg/mL, compared to the value of 2820 pg/mL of uninfected animals (p=0.0093), as shown in **Fig 5B**. Taken together, these results clearly indicated that activation of neutrophils/ NETs formation and platelet/endothelium had occurred at least at 4 d.p.i. in SARS-CoV-2-infected AC70 hACE2 Tg mice. Subsequent immunohistochemistry (IHC) staining of lung tissues derived from infected and uninfected AC70 hACE2 Tg mice with antibodies specific to platelet factor 4 (PF4), a specific biomarker of activated platelets, revealed extensive activation of platelets. Positive staining of PF4 was particularly intense around the areas showing profound blockages of veins and microcapillaries within the alveolar walls of the infected lungs (**Fig 5C**), as compared to non***-***infected tissues where only very few areas of weak staining of PF4 were detected (**Fig 5D**).

**Fig 5.**
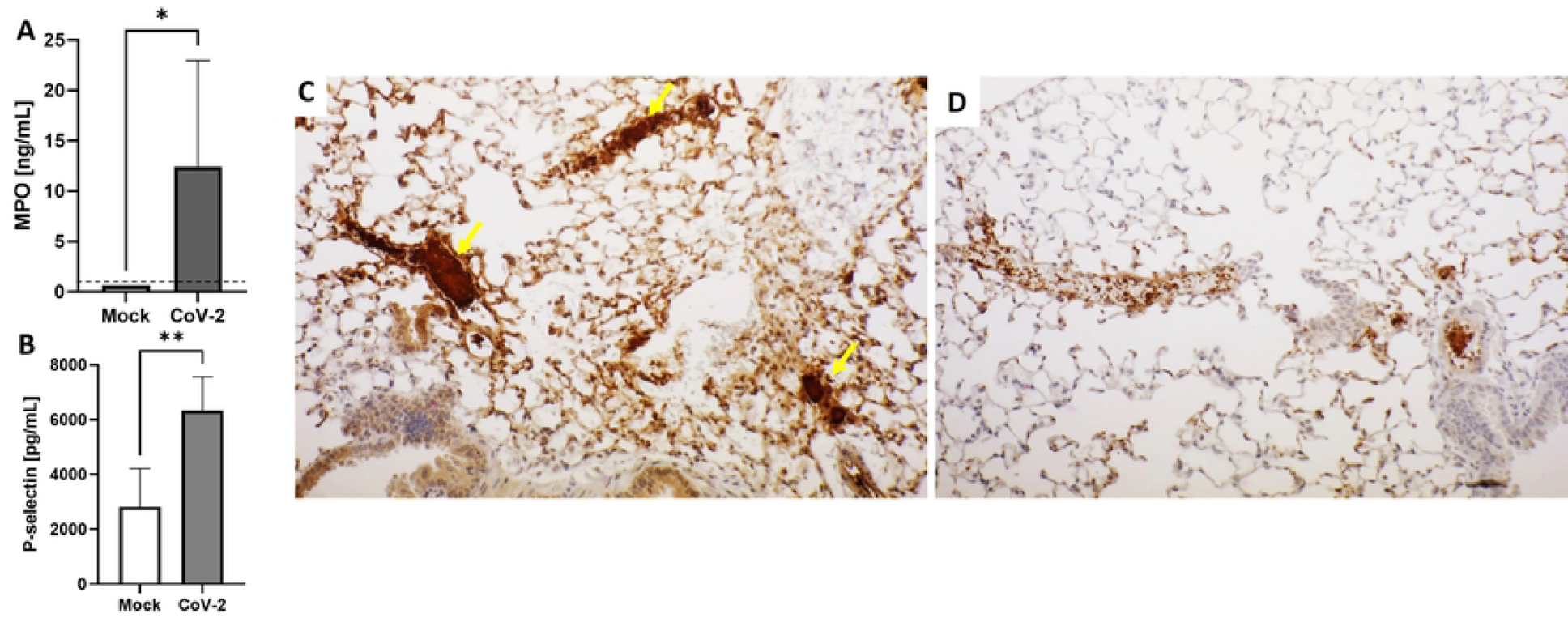
Lethal SARS-CoV-2 infection induced neutrophil/NETs and platelet/endothelium activation in AC70 hACE2 Tg mice. Evaluation of neutrophil/NETs and platelet/endothelium activation (**A&B**). Quantification of myeloperoxidase (MPO) (**A**) and P-selectin (**B**). Gamma irradiated EDTA-plasma samples collected from Mock mice and 4 d.p.i. with 10^5^ TCID_50_ of SARS-CoV-2 were analyzed by ELISA assays. Data are presented as means ± SEM (n=5 per group). Limit of detection for MPO assay was 0.61 ng/mL. Results were considered significant with p < 0.05 (*p=0.037, **p=0.0094). Representative immunohistochemistry (IHC) staining of lungs to Platelet Factor-4 (PF4) (**C&D**). Activation of platelets is shown as brown staining. Completely blocked veins and capillaries in the alveolar walls of the infected lungs (C, **yellow arrows**) were seen compared to the non-infected tissues (**D**). Bar scale: 80µm.

### Lethal SARS-CoV-2 infection induced autoantibody response to Annexin A2 in AC70 hACE2 Tg mice

Hospitalized COVID-19 patients that suffered from complications of coagulation and multiple organ failures had elevated autoantibody responses to various molecules, including phospholipids and phospholipid-binding proteins (aPL) [10]. Interestingly, levels of one of these antibodies targeting Annexin A2 (ANXA2) have been readily elevated among hospitalized patients who died from COVID-19 [11]. Acting as a co-receptor for tissue plasminogen activator, ANXA2 plays a critical role in the process of fibrinolysis at the cell surface. Thus, an elevated antibody response to ANXA2 in severe COVID-19 patients might interfere with the fibrinolytic process, causing undesirable thrombosis. We also investigated whether anti-ANXA2 antibodies could be detected in SARS-CoV-2-infected Tg mice by using an in-house developed ELISA assay. As shown in **Figure 6A**, we found that the levels of anti-ANXA2 antibodies in the circulation were indeed significantly increased at 4 d.p.i., compared to those of uninfected control (p=0036).

**Fig 6.**
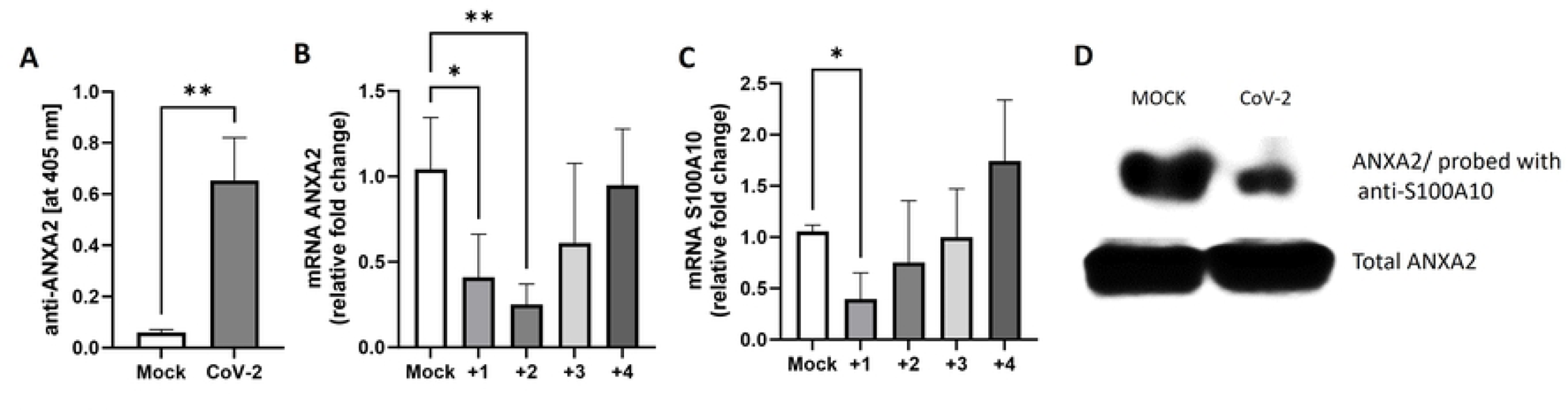
Lethal SARS-CoV-2 infection induced autoantibody to ANXA2 production and dysregulated ANXA2 system in AC70 hACE2 Tg mice. (**A**) Gamma irradiated EDTA-plasma samples collected from Mock mice and at 4 d.p.i. with 10^5^ TCID_50_ of SARS-CoV-2 were analyzed by ELISA assay. The results were expressed as absorbance values at 405 nm. The levels of (**B**) ANXA2 and (**C**) S100A10 gene expressions in lungs harvested from Mock and infected mice over the course of disease measured by RT-qPCR. The relative amount of targeted mRNA was obtained by normalizing with endogenous control gene (18S rRNA) and calculated in terms of the fold difference by the standard threshold cycle (ΔΔ*C_T_*) method. Data are presented as means ± SEM (n=5 per group). Results were considered significant with p < 0.05 (*p<0.05, **p<0.005). (**D**) WB shows decreased level of associated ANXA2 in S100A10 precipitates from lung tissue harvested at 4 d.p.i. compared to Mock specimens.

### SARS-CoV-2 infection downregulated transcriptional expressions of Annexin A2 and S100A10, and reduced the subsequent AIIt complex formation in AC70 hACE2 Tg mice

In addition to increasing the production of autoantibodies against ANXA2, we investigated if SARS-CoV-2 infection could dysregulate the transcription of the ANXA2-encoding gene. Specifically, we performed RT-qPCR to measure the transcripts of ANXA2 in the lungs daily of Mock-versus-SARS-CoV-2-infected mice. As shown in **Figure 6B**, we found that the levels of ANXA2 mRNA were promptly and significantly reduced to merely 41% (p=0.016) and 25% (p=0.0026) of that of uninfected control at 1 and 2 d.p.i., respectively, before gradual recovery thereafter. ANXA2 can exist as a monomer in cytoplasm or form heterotertrameric complexes (AIIt) with two S100A10 (p11) molecules at the extracellular surfaces. Thus, we also measured the mRNA levels of S100A10 within the lungs and found that like ANXA2, transcriptional expression of S100A10 was significantly downregulated to 39% (p=0.042) at 1 d.p.i., when compared to that of uninfected control, and gradually rebounded (**Figure 6C**). Taken together, these results indicate that SARS-CoV-2 infection effectively downregulated *de novo* transcription of both ANXA2 and S100A10 early during infection.

Since anti-ANXA2 antibody levels were significantly elevated at the later stage of SARS-CoV-2 infection in AC70 hACE2 Tg mice, we finally performed co-immunoprecipitation studies in lung tissues to determine whether the association between ANXA2 and S100A10 could be affected. As shown in **Figure 6D**, at 4 days after acute SARS-CoV-2 infection, the level of associated ANXA2 in S100A10 precipitates was readily decreased suggesting reduction of AIIt complex formation. Together, our data indicate that acute SARS-CoV-2 infection could modulate the ANXA2 system, thereby initiating cascades of impairment of ANXA2-dependent membrane-associated fibrinolysis in AC70 hACE2 Tg mice.

## Discussion

Over the past three years of the COVID-19 pandemic, various strategies have been employed to generate animal models for SARS-CoV-2, including non-human primates [26,27] ferrets [28], minks [29], hamsters [30, 31], and several mouse models, such as wild-type mice/mouse-adapted (MA) SARS-CoV-2 and various hACE2 transgenic (Tg) mice/clinical isolates SARS-CoV-2 models [32–37]. Even though proposed animal models mimic different clinical aspects and features of SARS-CoV-2 infection, none of them could fully recapitulate all aspects of COVID-19 observed in patients. Therefore, continuing effort to develop and characterize of animal models remains needed to better address the diverse spectrum of SARS-CoV-2 pathogenesis. Since coagulation disorders may be a key player in the onset of post-acute sequelae of COVID-19 [4], an animal model which could mimic aspects of coagulopathy as closely as possible to human patients is highly desirable to better understand the pathogenetic mechanisms of this phenomenon. Among these SARS-CoV-2 permissive animal species, mouse is one of the most prominent and the best-established species due to its availability, affordability, rich collection of immunological reagents and flexibility for various scientific purposes. We have previously developed transgenic AC70 hACE2 Tg mice that express the hACE2 gene under the synthetic CAG composite (CAG-hACE2) promoter for SARS-CoV infection showing weight loss and other clinical signs with 100% mortality within 8 days post-challenge [15, 16]. Here, we fully characterized the permissiveness of AC70 hACE2 Tg mice to SARS-CoV-2 infection and disease, including morbidity (weight loss and other signs of illness) and mortality, tissues tropisms and histopathology of viral infection, host inflammatory responses after a lethal dose of infection. These data along with well-determined values of LD_50_ and ID_50_ as low as ∼ 3 and 0.5 TCID_50_ of SARS-CoV-2, respectively, indicated that AC70 hACE2 Tg mice are highly permissive to SARS-CoV-2 infection, making it an additional useful model for *in vivo* evaluation of therapeutics and prophylactics against COVID-19 [38]. Additionally, we showed that lethally challenged AC70 hACE2 Tg mice manifested several hematological and coagulation abnormalities like those observed in clinical patients with PASC. We have also demonstrated that the fine balance between coagulation and fibrinolysis could be readily impaired and identified a possible role of Annexin A2 in coagulopathy developed after SARS-CoV-2 infection.

Intranasal challenge of AC70 hACE2 Tg mice with high dose of SARS-CoV-2 resulted in acute infection. Infected mice rapidly succumbed to disease, evidenced by body weight loss and development of clinical symptoms as early as 3 d.p.i. and eventually died by 5 d.p.i. We have assessed that the value of LD_50_ was only ∼3 TCID_50_, rendering AC70 hACE2 Tg mice more sensitive to SARS-CoV-2 compared to other Tg mouse models [36]. High levels of infectious virus were found in lungs and brains; however, viral RNA was detected in other tested tissues. Particularly, relatively high viral load was present in hearts, and noticeable in livers, spleens, kidneys, and GI tracks, indicating that lung and brain are the major target of SARS-CoV-2 while suggesting the disseminating nature of infection. The main pathological findings were seen in lungs and brains, however, expression of several pro-inflammatory cytokines and chemokines was also enhanced in other tested organs suggesting systemic inflammation.

Post-COVID complications are often observed in patients severely affected by SARS-CoV-2 infection and disease. Changes in various hematological parameters, particularly lymphopenia, and an elevated NLR value, are indicators of the systemic inflammatory response and have been used as predictors of severe viral or microbial infection and disease, including COVID-19 [39, 40]. Acutely decreased leukocytes, especially lymphocyte counts, concomitant with increased NLR value in AC70 hACE2 Tg mice were unambiguously associated with SARS-CoV-2 infection. The reason why lymphopenia is associated with severe illness remains unclear. It has been hypothesized that this association could be caused directly by virus attachment, indirectly by inflammatory mediators or infiltration of circulating lymphocytes into inflamed tissues [41]. Since neutrophils are the most abundant leukocytes in human circulation, leukocytosis associated with high neutrophil counts was also seen in patients. The balance of lymphocytes and neutrophils in mice is quite different with a strong preponderance of lymphocytes. This could explain leukopenia observed in infected AC70 hACE2 Tg mice, even though transient neutrophilia also occurred.

The role of red blood cells (RBC) in COVID-19 remains unknown. However, it has been suggested that SARS-CoV-2 infection has a negative impact on RBC parameters, resulting in the onset of anemia especially in severe patients with confirmed SARS-CoV-2 infection [42]. It has been reported that with aggravation of the disease, the HCT level and the concentration of Hb tend to progressively decrease; however, more exploratory studies are needed on this aspect. While we observed an inverse correlation in our AC70 hACE2 Tg mice at the peak of viral infection, additional studies are warranted. However, possible dehydration due to reduced water intake as the mice developed acute wasting syndrome, might explain this finding.

Although significant thrombocytopenia is common in many viral illnesses [43, 44], it is not as markedly observed in COVID-19. Mild thrombocytopenia has been variably observed in up to one-third of COVID-19 patients, however, thrombocytosis was reported in more severe disease as well [45–47]. We observed increased platelet counts at 3 d.p.i. in AC70 hACE2 Tg mice. Current reports indicate that COVID-related coagulopathy is characterized by thrombocytosis at the early stage and thrombocytopenia in the late phase of these conditions [22, 48].

The pathophysiology of COVID-19-associated coagulopathy is complex and may have a unique pattern compared to the standard mechanisms of thrombosis reported in other severe viral respiratory pneumonias. Our results indicate that AC70 hACE2 Tg mice had an abnormal coagulation and fibrinolytic system upon SARS-CoV-2 infection, as evidenced by markedly elevated D-dimer levels in the circulation. D-dimer is generated by the protease plasmin cleavage of cross-linked fibrin, an essential component of blood clots. Formation of cross-linked fibrin networks, and their subsequent enzymatic dissolution into D-dimer and other fibrin degradation products before leaking into the bloodstream might contribute to the elevated D-dimer levels in the plasma. Therefore, elevated levels of D-dimer in circulation might indicate the activation of both coagulable and fibrinolytic states. Fibrinolysis is tightly regulated by plasminogen activators, particularly t-PA, and inhibitors, such as PAI-1, with the conversion of plasminogen to proteolytically active plasmin that supports this process. In AC70 hACE2 Tg mice, although the presence of D-dimer along with the elevated expression of t-PA may at least indicate that fibrinolytic pathway remains functional to certain extents, concomitantly increased expression of PAI-1 levels suggests impaired fibrinolysis had occurred. The reason why fibrinolysis failure is observed despite high levels of t-PA remains unclear. One of the possible mechanisms is that high levels of PAI-1 overcome the local effect of t-PA producing a net hypofibrinolytic/ prothrombotic state in COVID-19 [49]. Another possible hypothesis describes the “plasmin paradox” in which fibrinolysis inhibition may occur at the early stage of COVID-19 whereas plasminogen activators accelerate fibrinolysis at later stages of disease [50].

The major source of t-PA and PAI-1 is likely derived from activated endothelium and/or release from activated platelets. It was suggested that extensive release of proinflammatory cytokines induced by the formation of neutrophil extracellular traps (NETs) triggers endothelial cell activation [51], thereby promoting local release of t-PA and PAI-1. Importantly, activated platelets within the microvasculature are deeply involved in neutrophil activation [2], which could lead to promotion of a hypercoagulation state within inflamed tissues. The ability to NET formation was also validated in SARS-CoV-infected AC70 hACE2 Tg mice. We demonstrated increased levels of both activated neutrophil/NET and platelet/endothelial cell markers in peripheral blood collected from SARS-CoV-2 infected mice, compared to non-infected animals. Additionally, mouse necropsies revealed extensive platelet activation within lung tissues with areas of platelet clumps that seem to block alveolar vessels. These findings are in line with the hypothesis that platelet and neutrophil activation along with endothelial dysfunction can contribute to tissue damage and widespread thrombosis in COVID-19.

Analysis of plasma samples collected from AC70 hACE2 Tg mice showed markedly elevated levels of IgG anti-ANXA2 antibodies, compared to those collected prior to infection. Autoantibodies targeting ANXA2 have been previously found in antiphospholipid syndrome (APS). Furthermore, their role in thrombotic events was demonstrated in association with APS [52]. Additionally, prior studies on sera collected from SARS-CoV patients revealed the presence of antibodies that reacted with ANXA2 on lung epithelial cells [13]. Importantly, a recent study reported elevated levels of anti-ANXA2 antibodies among hospitalized COVID-19 patients who died when compared to those of non-critically affected patients [11]. However, the importance of these autoantibodies remains unclear. Our data indicate that the induction of anti-ANXA2 occurred as early as 4 d.p.i. Rapid induction of autoantibodies targeting several autoantigens has been described in acute respiratory distress syndrome (ARDS) and severe patients with sepsis. It is intriguing that critically ill patients often displayed high titers of autoantibody within the first few days of admission to the Intensive Care Unit (ICU) [52]. Based on the fact that anti-ANXA2 detected in lethally challenged AC70 hACE2 Tg mice were of the IgG isotype, the typical phenotype of the adaptive immune response to high dose of SARS-CoV-2 challenge dose, is rather unlikely for this phenomenon. Thus, the expansion and differentiation of existing long-term memory B-cells specifically against ANXA2 or other autoantigens or inflammatory mediators might have been induced upon SARS-CoV-2 infection.

ANXA2 is a key player in cell surface fibrinolysis by acting as a co-receptor that orchestrates activation of endogenous tissue plasminogen activator tightly involved in fibrin clearance [12]. Therefore, dysregulation of ANXA2 mechanism may contribute to thrombotic events observed in COVID-19. We determined the kinetics of ANXA2 gene expression in lungs of AC70 hACE2 Tg mice by measuring mRNA levels over the time course of 4 d.p.i. and showed that transcriptional expression of ANXA2 was significantly downregulated between 1-2 d.p.i. The fibrinolytic potential of ANXA2 and its dysregulation have been well documented *in vivo* and *in vitro*. ANXA2-null mice (AnxA2^-/-^) manifested increased fibrin depositions in multiple organs, including lung, heart, and kidney compared to wild type mice, suggesting its importance in maintaining a balanced hemostasis [54]. The opposite effect, termed “annexinopathy”, has been described in acute promyelocytic leukemia (APL), in which overexpression of surface ANXA2 led to increased plasmin generation and hyperfibrinolysis [55]. On the cell surface ANXA2 is presented mostly as a heterotertameric complex with S100A10 molecules (AIIt). Interestingly, extensive *in vitro* and *in vivo* studies demonstrated a well-regulated collaboration of both molecules in t-PA-dependent fibrinolysis. Specifically, *in vitro* downregulation of ANXA2 resulted in reduced levels of S100A10 [56]. Additionally, it was noted in AnxA2^-/-^ mice that the expression of S100A10 was concomitantly depleted as well [57]. Furthermore, increased fibrin accumulation was also observed in tissues collected from S100A10-null mice [58]. Taken together, these reports strongly suggested that interaction of both proteins, in which ANXA2 functions by stabilizing S100A10 on the cell surface, is needed for surface fibrinolysis and plasmin generation. Thus, we measured S100A10 mRNA levels in lung specimens showing that gene expression was downregulated after SARS-CoV-2 infection with similar kinetics as ANXA2. Given these findings, decreased expression of both ANXA2 and S100A10 may contribute to the hypercoagulable state observed in AC70 hACE2 Tg mice. Interestingly, at the later phase of SARS-CoV-2 infection in our mouse model, i.e., 2 to 4 d.p.i., the expressions of ANXA2 and S100A10 rebounded to normal levels, suggesting that ANXA2/S100A10 may play a role in fibrinolytic failure at the early stage of acute infection. As mentioned above, the exact role of autoantibodies to ANXA2 in the COVID pathogenesis remains unknown and is currently under further investigation. We demonstrated that association between ANXA2 and S100A10 was reduced at 4 d.p.i., the time point when anti-ANXA2 antibodies were presented. Cesarman-Maus et al. [52] raised the hypothesis that anti-ANXA2 autoantibodies may promote hypercoagulation and fibrin deposition by inhibiting t-PA-dependent plasmin formation. This led us to hypothesize that during acute SARS-CoV-2 infection the levels of t-PA may remain sufficient to sustain an effective fibrinolysis, however, the induction of anti-ANXA2 antibodies might negatively affect t-PA binding by disrupting the formation of AIIt complex.

In summary, we have characterized acute SARS-CoV-2 infection and post-acute coagulation sequalae associated with COVID-19 in AC70 hACE2 Tg mice that are proven highly permissive to SARS-CoV-2 infection and disease. Since disease characteristics of infected AC70 hACE2 Tg mice recapitulates many of the pathological findings observed in human patients, these mice can be used not only for translational research but offers an attractive animal model for studying molecular and cellular mechanisms involved in the pathophysiology of COVID-associated coagulopathy. Taking advantage of AC70 hACE2 Tg mice we highlighted the potential role of Annexin A2 in this process, which can serve as a promising strategy to reduce thrombotic events that frequently occurring in severe disease. Additional studies are warranted to further explore the impact of ANXA2 mechanism and identify potential therapeutic targets for COVID-19.

## Materials and Methods

### Ethics statement

All experiments involving animal and infectious virus were carried out at the Galveston National Laboratory at The University of Texas Medical Branch at Galveston (UTMB), Texas, an AAALAC accredited (November 24, 2020) and PHS OLAW approved (February 26, 2021) high-containment National Laboratory, based on a protocol approved by the Institutional Animal Care and Use Committee (IACUC) at UTMB Galveston.

### Virus stock and cells

SARS-CoV-2 (US_WA-1/2020 isolate), provided through the World Reference Center for Emerging Viruses and Arboviruses (WRCEVA), was used throughout the study. Viral stock was expanded in Vero E6 cells (American Type Culture Collection) by passaging three times and resulting cell-free supernatants were sequenced, titrated, and stored at – 80°C as working stocks.

### Mice challenge and monitoring

Human ACE2 transgenic mice, line AC70, and their non-transgenic littermates were obtained from Taconic Biosciences (NY, Cat # 18222). Six-to-eight-week-old mice were anaesthetized with isoflurane in oxygen and inoculated intranasally (IN) with 10^5^ TCID_50_ of SARS-CoV-2 in 60 μl of Eagle’s minimum essential medium (EMEM) supplemented with 2% heat-inactivated Fetal Bovine Serum (FBS) (M-2). Infected animals were monitored at least once daily for weight changes, other signs of illness, and mortality.

### Tissue and blood collection

Blood samples were collected via retroorbital (RO) route under anesthesia to analyze the complete blood count (CBC) using a VetScan analyzer (VSpro, ABAXIS) and coagulation parameters using ELISA assays. At the indicated time point mice were euthanized using CO2 inhalation and several tissue specimens were collected. Briefly, the right lung was washed with Phosphate Buffered Saline (PBS) and inflated with 10% neutral buffered formalin for histopathology and immunohistochemistry. The left lung and other tissues were divided for downstream experiments, including viral titration, RNA and protein extraction.

### Viral titration

For quantifying infectious virus, tissues collected were weighed and homogenized in PBS/10% FBS using the TissueLyser (Qiagen; Retsch, Haan, Germany). After clarification by centrifugation, supernatants were titrated in Vero E6 cell cultures grown in 96-well microtiter plates using a standard infectivity assay [15]. Viral titers in homogenates were expressed as TCID_50_/g of tissue (log_10_).

### Histopathology (H&E) and immunohistochemistry (IHC)

For H&E staining, pieces of tissues were fixed in 10% neutral buffered formalin (Sigma) for 72 hours, transferred to 70% ethanol and paraffin embedded. Staining on deparaffinized sections was performed by using routine hematoxylin-and-eosin method. IHC assessment following the previously published protocol [15]. CXCL4/PF4 (6Q10H2) (Cat# NBP3-16251) rabbit monoclonal antibodies were purposed from Novus Biologicals.

### ELISA assay

Gamma-irradiated EDTA-plasma samples collected from infected and uninfected mice were used for ELISA assays to detect components of the coagulation system. Commercially available kits were used for assessment of the levels of Myeloperoxidase (Cat #EMMPO) and PAI-1 (Cat# EMSERPINE1) purchased from Invitrogen, TPA (Cat# ab233615) and P-Selectin (CD62P) (Cat# ab200014) purchased from Abcam, and D-Dimer (D2D) (Cat# RK02735) purchased from ABclonal. Plates were read using an ELISA plate reader (Molecular Device). To assess the levels of anti-Annexin A2 (ANXA2) antibodies, 200ng of recombinant ANXA2 protein (Cat# 9409-AN, R&D Systems) diluted in PBS, pH 7.4, was used to coat an Immulon 2 HB 96 well-plate (Thermo Fischer) overnight at 4°C. Plates were washed 3 times with PBS/ 0.1% Tween-20, pH 7.4 (PBS-T) using a Microplate 405 washer (BioTek). 100 μl plasma samples diluted 1:10 in plasma diluent (PBS/5% skim milk/0.1% Tween-20) were inclubated for 2 hours at 37°C. After the incubation, plates were washed 3 times with PBS-T. 100 μl anti-mouse IgG antibodies conjugated to the enzyme horseradish peroxidase (HRP) (Invitrogen) diluted at 1:2000 in plasma diluent were added and incubated at 37°C for 1 hour. After the final washing step, plates were incubated with ABTS substrate (Thermo Fisher) for 30 minutes at 37°C. Plates were read at 405 nm using a PerkinElmer Victor XV plate reader.

### Co-immunoprecipitation (IP) and Western Blot (WB)

Gamma-irradiated infected and non-infected tissue specimens were homogenized on ice using Dounce Glass Manual Tissue Grinder (Etcon) following the Plasma Membrane Protein Extraction Kit protocol (Cat# ab65400, Abcam). Total protein concentration was quantified using Pierce™ BCA Protein Assay Kit (Thermo Fisher). For IP, anti-S100A10 antibody [EPR3317] (Cat# ab76472) (Abcam) were immobilized onto an AminoLink Plus Coupling Resin (Thermo Fisher) and incubated for 2 hours at room temperature. Equal amounts of tissue lysates were pre-cleared using the Control Agarose Resin (Thermo Fisher). The antibody-antigen complexes were incubated overnight at 4°C and eluted following Pierce Co-Immunoprecipitation (Co-IP) Kit (Thermo Fisher). Samples were then separated by gel electrophoresis followed by immunoblotting. For the WB, equal amounts of immunocomplexes were subjected to 4–20% SDS-polyacrylamide gel (Bio-Rad) electrophoresis (SDS-PAGE). Proteins were transferred onto a polyvinylidene difluoride membrane and then incubated with anti-annexin A2 (ANXA2) rabbit polyclonal antibodies (Cat# ab41803), followed by incubation with Eu-Labeled Goat Anti-Rabbit ScanLater for 1 hour. Blots were visualized using ScanLater Western Blot Detection System (Molecular Devices).

### RNA extraction and real-time quantitative qRT-PCR

Pieces of tissues were transferred to individual vials containing RNAlater solution (Qiagen) and subsequently homogenized as described above. Total RNA was isolated using TRIzol (Life Technologies), treated with Turbo DNAse I (Invitrogen) and quantitated using SpectraMax Paradigm Microplate Reader (Molecular Device). Complementary DNA was prepared with iScript Reverse Transcription Supermix for qRT-PCR (Bio-Rad), and qRT-PCR was carried out with specific primers (**Table S2**) and iQ SYBR green Supermix (Biorad) using CFX96 real time system (Biorad). The relative amount of targeted mRNA was obtained by normalizing with endogenous control gene (18S rRNA) and calculated in terms of the fold difference by the standard threshold cycle (ΔΔ*C_T_*) method [59].

### Statistical Analysis

Statistical analyses were performed using GraphPad Prism software. Values from multiple experiments are expressed as means ± SEM. Statistical significance was determined for multiple comparisons using one-way analysis of variance (ANOVA). Student’s t-test was used for comparisons of two groups. Results were regarded as significant if two-tailed P values were < 0.05. All data are expressed as mean ± standard error of the mean.

## Acknowledgments

We thank Dr. Pinghan Huang for his technical support. We are grateful to members of the Animal Resources Center for their help with the logistics and providing quality care for research animals, and the Research Histology Core for tissue processing. We thank Dr. Kenneth S. Plante for providing the virus strain, and the World Reference Center for Emerging Viruses and Arboviruses (WRCEVA), Dr. Natalie Thornburg and the Centers for Disease Control and Prevention (CDC) for graciously donating the virus to the collection. We also thank Drs. Eric D. Lazartigues and Abraam Yakoub for their editorial review of the manuscript.

## Supporting information

**Fig S1. Cytokine and chemokine responses to lethal SARS-CoV-2 infection in AC70 hACE2 Tg mice.** AC70 Tg⁺ mice were challenged intranasally with 10^5^ TCID50 of SARS-CoV-2 and sacrificed at 4 d.p.i. Total RNA was isolated from (**A**) lungs, (**B**) brains, (**C**) spleens, (**D**) livers, and (**E**) kidneys, and cytokines and chemokines expressions were quantified by RT-qPCR. Results are shown as mean relative fold change in expression compared to Tg^-^ mice and were normalized to 18S RNA. *p < 0.05, **p < 0.01, ***p < 0.001 Tg⁺AC70 vs Tg^⁻^.

**Fig S2. Determining the 50% lethal dose (LD_50_) of SARS-CoV-2 (US_WA-1/2020) in AC70 hACE2 Tg mice.** Mice (n=6) were challenged with 1 × 10^1^, 1 × 10^0^ and 1 × 10^-1^ TCID_50_ of SARS-CoV-2. (**A**) % body weight change, and (**B) %** survival. Estimated LD_50_ is ∼3.

**Table S1. Determining the 50% infectious dose (ID_50_) of SARS-CoV-2 in AC70 hACE2 Tg mice.** SARS-CoV-2 antibody response was determined by analysis of serum specimens from survived mice at 21 d.p.i. ND: not detected; NA: not applicable. Estimated ID_50_ is ∼0.51 TCID_50._

**Table S2. The primer sequences.**

